# Antisense oligonucleotide to SARS-Cov-2 trs gene: antiviral activity on an in vitro model and possibilities of vital and postmortem diagnosis of COVID-19

**DOI:** 10.1101/2022.08.13.503857

**Authors:** A.N. Goryachev, A.A. Shtro, A.V. Galochkina, A.S. Goryacheva

## Abstract

The data on the relevance of the 5 ‘-AGC CGA GTG ACA GCC ACA CAG antisense oligonucleotide for binding to the trs-gene of the SARS-CoV-2 virus, which causes the new coronavirus infection COVID-19, are presented. The high stability and conservatism of this section of the SARS-CoV-2 genome is shown, which allows it to be used as an application point for antisense oligonucleotides. By evaluating plaque inhibition, the ability of this antisense oligonucleotide with phosphorothioate and 2’-oxymethyl modification to suppress viral replication was found. The effective dosage reducing the virus titer by 50% is 3.84 mcg/ml. No toxicity was shown up to a dosage of 100 μg/mL, which is more than 28.8 chemotherapeutic index. The ability of this oligonucleotide conjugated to the fluorescent dye TAMRA to detect the SARS-CoV-2 virus in the fluorescent hybridization reaction in situ in cytological preparations of nasopharyngeal smears and blood smears, as well as in histological preparations of internal tissues is shown.

## RELEVANCE

Over the past three years, the number of biological threats in the world has increased. First of all, these are viral diseases such as COVID-19, monkey smallpox virus, Marburg hemorrhagic fevers and others. The example of COVID-19 shows that epidemic viral diseases have a multiplicative effect, causing growing problems not only in the healthcare sector, but also in the economy as a whole.

Such challenges to society require modern high-tech solutions based on knowledge of the mechanisms of the spread of viruses. One such solution for viral diseases is the use of antisense oligonucleotides (ASO). ASO are single-stranded oligonucleotide strands of DNA or RNA from 20-25 nucleotides, complementary to genomic viral RNA sequences (10).

The feasibility of using ASO for the treatment of Coronaviridae family viruses (SARS-Cov-Tor, MERS) has previously been confirmed by studies (8). At the same time, the most optimal is the use of ASO to the initial sections of the coronavirus genome, in the zone of the gene regulating transcription (trs), which is located near the 5 ‘-end of the beta-coronavirus genome. Specific binding of ASO to this génom leads to blocking the transcription of almost the entire viral genome, which prevents further reproduction of the virus and the synthesis of viral proteins.

We previously studied ASO of the type 5 ‘-AGC CGA GTG ACA GCC ACA CAG with phosphorothioate modification of the internucleoside bond, complementary to the section of the trs-gene of the SARS-Cov-2 coronavirus. A study of the antiviral activity of this ASO showed a decrease in virus replication by 9-13 times (3). However, the emergence of new strains of coronavirus dictates the need to study the antiviral activity of new ASO or modify existing ASO for the treatment of a new coronavirus infection (NCI) COVID-19.

In addition, highly specific binding of selected ASO to conserved regions of the SARS-Cov-2 genome can be used to conjugate ASO data to fluorescent markers to detect viral RNA in an fluorescence in situ hybridization (FISH) reaction in cytological and histological studies (6). Moreover, these studies can be of value not only for intravital, but also for autopsy diagnostics, including in retrospective studies of autopsy material. Studies of viral RNA in tissues by the FISH method are known and used to reveal the mechanisms of pathogenesis of the new coronavirus infection COVID-19 (4, 12). In this regard, it seems relevant to study the antiviral activity of the known ASO to the SARS-Cov-2 of the 5 ‘-AGC CGA GTG ACA GCC ACA CAG species with additional modification, as well as the possibility of using this ASO for lifetime and post-mortem diagnosis of SARS-Cov-2 in the FISH response, which suggests the relevance of this study.

**THE PURPOSE** of the study is to determine the antiviral activity of the known ASO to the SARS-Cov-2 of the species 5 ‘-AGC CGA GTG ACA GCC ACA CAG with phosphorothioate modification of internucleoside phosphate and 2’ -oxymethyl modification of pentose, as well as to determine the possibility of using this oligonucleotide markered with a fluorescent marker for cytological and histological diagnosis of SARS-Cov-2. In accordance with the purpose, the objectives of the study were determined.

1. Based on BLAST analysis, investigate the relevance of the proposed ASO for all possible SARS-Cov-2 mutations.
2. To evaluate the antiviral activity of the proposed ASO with phosphorothioate modification of internucleoside phosphate and 2’-oxymethyl modification of pentose in vitro against SARS-CoV-2 coronavirus.
3. To investigate the applicability of the proposed ASO conjugated with the fluorescent tetramethylrodamine marker (TAMRA) at the 5’-end on cytological smears with nasopharyngeal mucosa and blood smears in patients with an established diagnosis of NCI COVID-19, as well as on histological sections of internal organ tissues of persons who died with an established diagnosis of NCI COVID-19.

## MATERIAL AND METHODS

1. A study of the relevance of the proposed ASO of type 5 ‘-AGC CGA GTG ACA GCC ACA CAG was carried out by BLAST analysis (https://blast.ncbi.nlm.nih.gov/Blast.cgi). The need to check the operability of the ASO using the *in silico* method was dictated by the high mutation potential of the coronavirus, as a result of which the target nucleotide sequences of the virus genome could change and lose relevance as a point of the ASO application. To block transcription, a section of the coronavirus gene responsible for transcription of trs was chosen. This site has the sequence 5 ‘-CTG TGT GGC TGT CAC TCG GCT, the presence of which was checked in the sequenced nucleotide sequences of all identified SARS-Cov-2 samples (SARS-CoV-2 Data Hub, NCBI) (9).
2. The synthesis of ASO to SARS-Cov-2 of the 5 ‘-AGC CGA GTG ACA GCC ACA CAG type with phosphorothioate modification of internucleoside phosphate and 2’-oxymethyl modification of pentose was carried out at DNK-Synthes LLC (https://oligos.ru/) in the amount of 4 mg (101 OE) in the classic amidophosphite method with further HPLC purification
3. The antiviral activity of ASO was tested by evaluating the inhibition of plaque formation (7) in the chemotherapy laboratory of viral infections of the A.A. Smorodintsev Influenza Research Institute of the Ministry of Health of Russia (head of the laboratory, Ph.D. A.A. Shtro; Research work LHT-SA-010/2022 from the 03.06.2022). Vero E-6 cell culture (kidney epithelium of green monkey) obtained from the working collection of the laboratory of chemotherapy of viral infections of the A.A. Smorodintsev Research Institute of Influenza was used for the study. Vero cells were cultured by intermittent cell transfer to T-75 vials (once every 3-4 days) using 1:1 Trypsin-Versen solution. After visual inspection of the cell slide, DMEM growth medium containing 10% fetal bovine serum (FBS) and antibiotic solution were added to the vial in such a volume that the resulting slurry contained about 3×106 cells/ml. For experimental work, cells were seeded into 6-well plates at an inoculated dose of 9×106 cells per well. Experiments were performed after complete confluence of the monolayer was formed, about 24 hours later. The work used clinical isolate No. 3524, obtained from the material of a patient with coronavirus infection by isolation in Vero E-6 cell culture, at the fifth passage in Vero cell culture. A sample of the virus was provided by the virological laboratory complex of the Institute of Experimental Medicine. Virus titration on Vero cell culture was performed on 6-well culture plates prior to experiment. A series of 10-fold consecutive dilutions was prepared from a sample of virus-containing suspension, each dilution was applied to a cell monolayer and incubated for 2 hours. Further, the virus-containing liquid was removed, after which the coating medium with 0.9% agar was applied and incubated for 3 days. At the end of the incubation period, the cells were fixed and stained with a 0.2% solution of crystalline violet in a 10% formaldehyde solution, after which the number of plaques formed was counted in each well. The smallest amount of virus tested leading to the formation of one plaque (lysis zone) is taken as the infectious unit (plaque-forming unit, PFU). SARS-Cov-2 virus effluent (hCoV-19/Russia/StPetersburg-RII3524VR4/2020 No. 35245) stored at -25 ° C for subsequent experiments was diluted to a titer of 100 TCID_50_ (50% tissue culture infective dose - a dose of virus that causes infection of 50% of cells), which corresponds to a value of 3*10 PFU. The experiment used a prophylactic regimen for the use of drugs 1 hour before infection with the virus. 6-well plates with a 100% monolayer of cells were used to study the activity of the preparations according to the prophylactic scheme. The supernatants were removed from the wells of the plates containing the cell monolayer immediately before the dilutions of the test object were introduced, after which 1 ml of 3-fold dilutions of the test objects in double concentrations were added to the corresponding wells of the plates and incubated with the cells for 1 hour at 37°C, 5% CO_2_. One well of the plate was used as an infection control - instead of diluting the test object, DMEM culture medium was added. Next, a 100 TCID_50_ viral inoculum was added to the wells of the plate and incubated for 2 hours at 37 ° C, 5% CO_2_. The culture medium (containing ASO and virus) was then removed, after which a coating medium containing 0.9% agar was added to the cells, to which ASO was also added in appropriate concentrations and further incubated for 3 days at 37 ° C and 5% CO_2_. At the end of the incubation period, the cells were fixed and stained with a 0.2% solution of crystalline violet in 10% formaldehyde, after which the number of plaques in each well was counted. In the experiments according to the prophylactic scheme, the following ASO concentrations were used − 100; 10; 1; 0.1; 0.01 mcg/ml. Virus titer (T) was calculated using the formula (1):

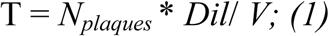

ГДe *N*_*plaques*_ – number of plaques, *Dil* – dilution of virus-containing material, *V* – the volume of the infectious concentration of the viral inoculum. Based on the obtained data, % inhibition of viral activity was calculated for each concentration of the test object, as well as ED_50_ - effective dosage, which caused a decrease in the virus titer by 50%.
4. The synthesis of the ASO fluorescent marker to the 5 ‘-AGC CGA GTG ACA GCC ACA CAG SARS-Cov-2 was carried out at DNK-Synthes LLC (https://oligos.ru/) in an amount of 50 OE by the classic amidophosphite method with further conjugation with tetramethylrodamine (TAMRA) for 5’-end and HPLC purification. Dilution of oligonucleotide in formamide at a concentration of about 100 ng/mcl (11) was used as the working concentration of the DNA probe
5. The study of the diagnostic effectiveness of the TAMRA-conjugated ASO was carried out by fluorescence hybridization in situ on cytological smears of the nasopharyngeal mucosa and blood smears in patients diagnosed with NCI COVID-19, as well as on histological sections of internal organs of individuals who died with an established diagnosis of NCI COVID-19. For the cytological study, nasopharyngeal mucosal smears and blood smears were taken from 14 patients. Patients by disease severity were distributed as follows. The control group was made up of healthy respondents aged 26 to 65 years in the amount of 8 people without clinical manifestations of NCI, with negative PCR results SARS-Cov-2 nasopharyngeal smears. Patients in the trial groups showed positive PCR results SARS-Cov-2 nasopharyngeal smears and different severity of NCI. Thus, the first experimental group included 5 people with an easy course of NCI. The second trial group included 6 patients with moderate NCI. The third trial group was 5 patients with a severe course of NCI. The severity of the disease was assessed in accordance with the Treatment Guidelines of National Institutes of Health (2). Cytological material was taken in the department for the treatment of patients with a NCI of the City Clinical Hospital of Pyatigorsk. Patients had nasopharyngeal smear and venous blood taken into EDTA tubes (purple cap). The collection of diagnostic material, its packaging, markering and transportation was carried out in accordance with the requirements and rules for working with materials potentially infected with pathogens of the II pathogenicity group, their storage and transportation in accordance with MU 1.3.2569-09 “Organization of the work of laboratories using NAAM (nucleic acid amplification methods) when working with material containing microorganisms of pathogenicity groups I-IV” and Methodological recommendations of Rospotrebnadzor 3.1.0169-20 “Laboratory diagnostics of COVID-19”. Smears were made on slides with Superfrost Plus (Thermo FS) adhesive coating. The smears were fixed in 100% ethanol (1). Next, blood and nasopharyngeal smears were prepared for fluorescent in situ hybridization (FISH) using a set of reagents for sample preparation of cytological preparations ("Medico-Diagnostic laboratory" LLC, www.m-d-l.ru). The preparations were treated with a solution of the SARS-Cov-2 ASO fluorescent marker in formamide. The centromeric DNA probe CCP1 FISH Probe, locus 1p11-q11 (CytoTest Inc.) was used as a probe control. The prepared preparations were examined under immersion on a fluorescent trinocular microscope Micromed 3 LUM with a green luminescent block (exciting radiation 500–550 nm, blocking light filter 590 nm). Microphotography was carried out with a Canon EOS 1100D camera. In parallel, standard staining of cytological preparations according to Romanovsky-Giemsa was carried out.
6. The applicability of the proposed ASO fluorescently markered tetramethylrhodamine (ASO-TAMRA) was investigated on histological sections of visceral tissues obtained at autopsy. Autopsy material was obtained in the Department of pathology of the City Clinical Hospital of Pyatigorsk. The experimental group of respondents consisted of 6 persons aged 58-72 years who died with an established diagnosis of NCI COVID-19. In all studied cases of death of persons in the experimental group, a severe life course of NCI was observed. The diagnosis of NCI COVID-19 was established antemortem according to a characteristic clinical picture, a positive SARS-Cov-2 PCR result and confirmed by post-mortem examination. The controls were 5 individuals aged 64–74 years with a long-term aggravated cardiac history, whose death occurred from acute coronary death within 24 hours from the onset of the attack, in whom, during intravital and postmortem SARS-Cov-2 PCR studies, negative, and there were no intravital clinical and post-mortem morphological signs of NCI. During autopsy, tissues of internal organs were taken, from which histological preparations were made according to the standard method with staining with hematoxylin and eosin (5). In parallel, for fluorescent hybridization, histological sections of tissues of internal organs were mounted on glasses with an adhesive coating Superfrost Plus (Thermo FS). Preparation for staining with the ASO-TAMRA fluorochrome probe was carried out using a set of reagents for sample preparation of paraffinized tissue sections ("Medico-Diagnostic laboratory" LLC, www.m-d-l.ru). The preparations were treated with a solution of the ASO-TAMRA fluorescent marker for SARS-Cov-2 with formamide. The centromeric DNA probe CCP1 FISH Probe, locus 1p11-q11 (CytoTest Inc.) was used as a probe control. The prepared preparations were examined under immersion on a fluorescent trinocular microscope Micromed 3 LUM with a green luminescent block (exciting radiation 500–550 nm, blocking light filter 590 nm). Microphotography was carried out with a Canon EOS 1100D camera.

## RESULTS OF THE RESEARCH

### I. Study of the relevance of the proposed ASO for all possible SARS-Cov-2 mutations

Checking the relevance of the proposed ASO species 5’-AGC CGA GTG ACA GCC ACA CAG by BLAST analysis showed that the complementary trs region of the 5’-CTGTGTGGCTGTCACTCGGCT gene is present only and exclusively in the SARS-Cov-2 genome. The presence of this region was verified in sequenced full genome (complete genome) nucleotide sequences of all identified SARS-Cov-2 samples (about 6 million records), where this fragment, according to the analysis of FASTA nucleotide sequences, was found in the region of the 5’-end of the genome at a distance of up to 84 nucleotides from the 5’ end of the chain.

### II. Study of the antiviral activity of ASO

The antiviral efficacy of the ASO preparation was evaluated against coronavirus by the method of plaque formation inhibition in various dosages according to the prophylactic scheme. The results are presented in tab 1.

**Table 1.** The results of the study of the antiviral activity of the ASO preparation by the method of plaque formation inhibition.

| ASO dose, mcg/ml | Virus titer, PFU |  |  | M±SD, PFU | Virus quantity, % |  |  | M±SD, % | ED<sub>50</sub> mcg/ml |
| --- | --- | --- | --- | --- | --- | --- | --- | --- | --- |
|  | 1 repeat | 2 repeat | 3 repeat |  | 1 repeat | 2 repeat | 3 repeat |  |  |
| 100 | 5,1<br>*10<sup>5</sup> | 5<br>*10<sup>5</sup> | 2<br>*10<sup>5</sup> | 4±1,8<br>*10<sup>5</sup> | 26,5 | 33,7 | 11,1 | 23,8<br>±11,5 | 3,84 |
| 10 | 9,6<br>*10<sup>5</sup> | 7,2<br>*10<sup>5</sup> | 6,6<br>*10<sup>5</sup> | 7,8±1,58<br>*10<sup>5</sup> | 50 | 48,6 | 36,6 | 45,1<br>±7,3 |  |
| 1 | 1,44<br>*10<sup>6</sup> | 1,28<br>*10<sup>6</sup> | 1,32<br>*10<sup>6</sup> | 1,3±0,8<br>*10<sup>6</sup> | 75 | 86,4 | 73,3 | 78,2<br>±7,1 |  |
| 0,1 | 1,88<br>*10<sup>6</sup> | 1,72<br>*10<sup>6</sup> | 1,68<br>*10<sup>6</sup> | 1,8±0,1<br>*10<sup>6</sup> | 97,9 | 116,2 | 93,3 | 102,4<br>±12,1 |  |
| 0,01 | 2,08<br>*10<sup>6</sup> | 1,52<br>*10<sup>6</sup> | 1,36<br>*10<sup>6</sup> | 1,7±0,37<br>*10<sup>6</sup> | 108,3 | 102,7 | 75,5 | 95,5<br>±17,5 |  |
| 0 | 1,92<br>*10<sup>6</sup> | 1,48<br>*10<sup>6</sup> | 1,8<br>*10<sup>6</sup> | 1,7±0,22<br>*10<sup>6</sup> | 100 | 100 | 100 | 100 |  |

From the above data, it can be seen that all the studied drug demonstrated the presence of antiviral activity in the studied concentrations. According to the results obtained, the drug had antiviral protection in the prophylactic regimen, which is confirmed by the low value of ED_50_ = 3.84 mcg/ml.

To visualize the results obtained, photographs of one of the experimental plates with negative viral colonies “plaques” for the experiments are shown. The data are shown in fig. 1.

**Fig. 1.**
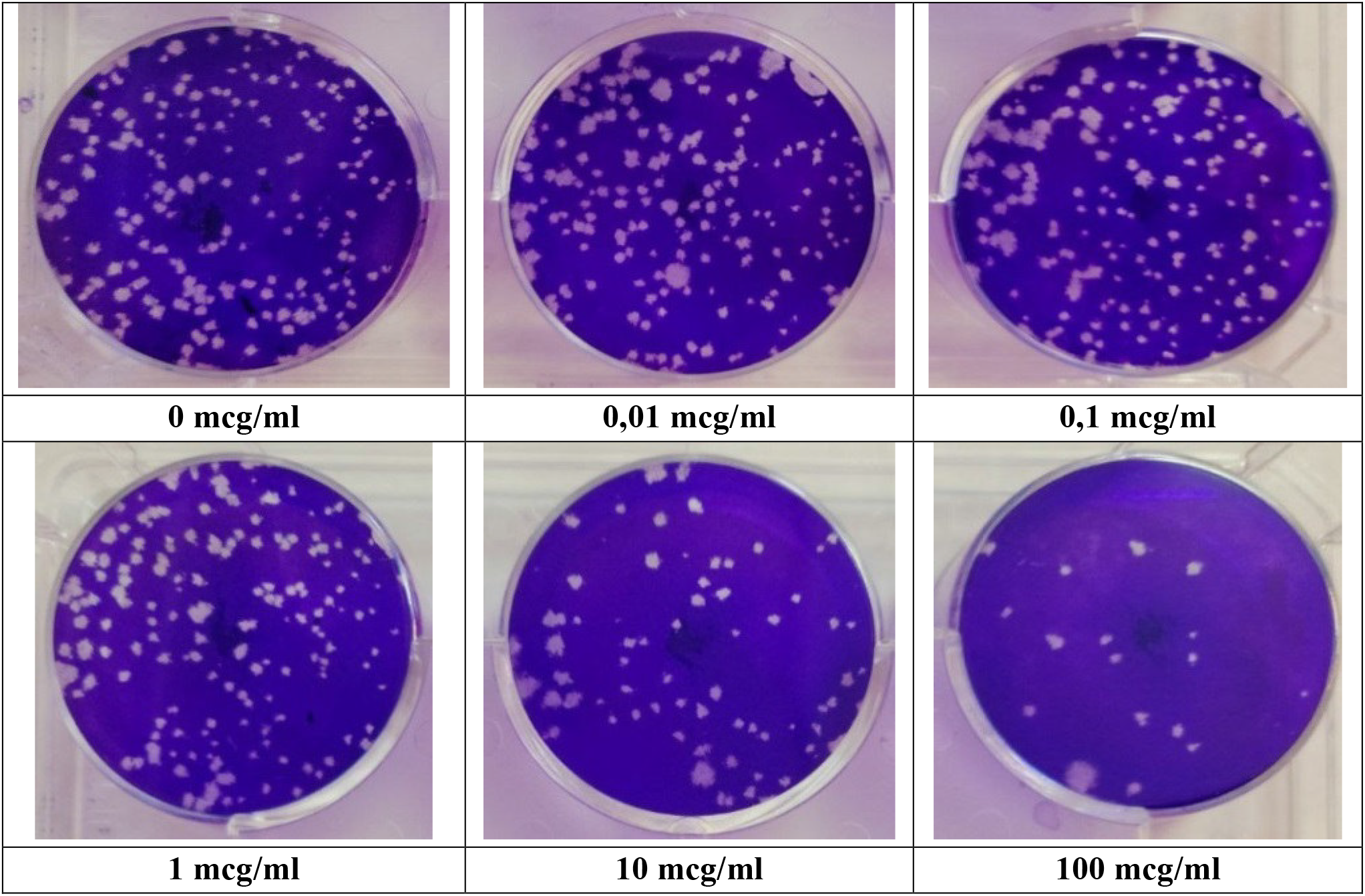
Results of testing the antiviral activity of the drug against the SARS-CoV-2 coronavirus.

Cytotoxicity testing of the drug was not carried out and the exact value of the cytotoxic dose that causes the death of 50% of cells - CTD_50_ cannot be calculated. However, to calculate the chemotherapeutic index (CTI), it is possible to use a concentration of 100 μg/ml, since no violations of the viability of the cell culture were noted during visual observation in the presence of the test drug at this concentration.

Thus, the value of the CTI of the drug is more than 100/3.48 = 28.8, which significantly exceeds the threshold value of 8, so it seems possible to conclude that the study drug has a high antiviral activity.

### III. Study of the diagnostic efficiency of ASO conjugated with TAMRA in the FISH reaction

Cytological examination of nasopharyngeal swabs in the FISH reaction revealed that in the control groups there was no fluorescence of the hybridized marker in the cells, which indicates the absence of viral particles. In mild NCI, a few epithelial cells were found containing single foci of the luminous ASO-TAMRA marker hybridized with SARS-Cov-2 RNA (Fig. 2 A, B). The number of virus-positive cells was 1-2 per field of view at small magnification of the microscope 10*10.

**Fig. 2.**
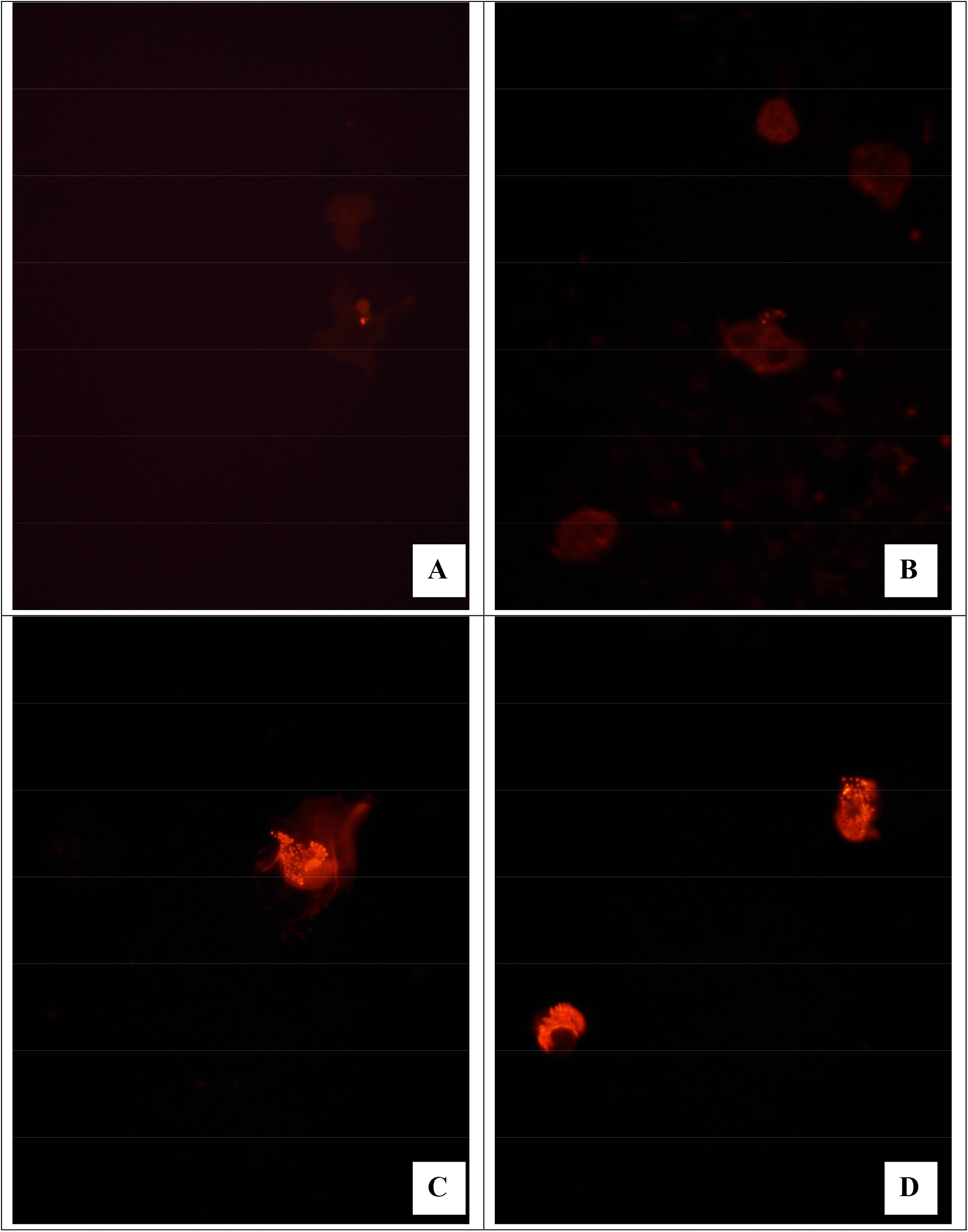
**FISH reaction to RNA SARS-Cov-2 in nasopharyngeal smears. Fluorescence microscopy. (A, B) - light course, single viral inclusions in the cytoplasm of the epithelium. (C) - is a moderate course. Abundance of viral RNAs in nasopharyngeal epithelial cells. (D) - is a severe course. Multiple cells with viral involvement. Multiplicity of magnification 10 * 100, immersion.**

In the case of a moderate course of NCI, multiple cells were detected (4-5 in the field of view at a small magnification of the microscope of 10*10), in which intensive reproduction of the virus in epithelial cells was detected. Moreover, the markers hybridized in coronavirus were located both in the nucleus and in the cytoplasm (Figure 2 C). ASO-TAMRA marks were detected in the amount of 10-15 field of view at a small magnification of the microscope of 10 * 10 in severe course of NCI. Virus-positive cells were also saturated with viral particles (Figure 2 D).

When examining blood smears, the cytological pattern was less pronounced. No ASO-TAMRA fluorescence was detected in the control group. No fluorescent markers were also found in venous blood cytological smear in individuals with mild NCI. Single cell elements (red blood cells) containing single viral RNA particles were found in patients of the experimental group with a moderate course of NCI (Figure 3A). The blood of patients with a severe course of NCI was more intense in the content of viral particles. In this group of patients, the fluorescence of ASO-TAMRA markers was detected in many fields of view, and the content of virus particles was sufficiently saturated (Fig. 3B).

**Fig. 3.**
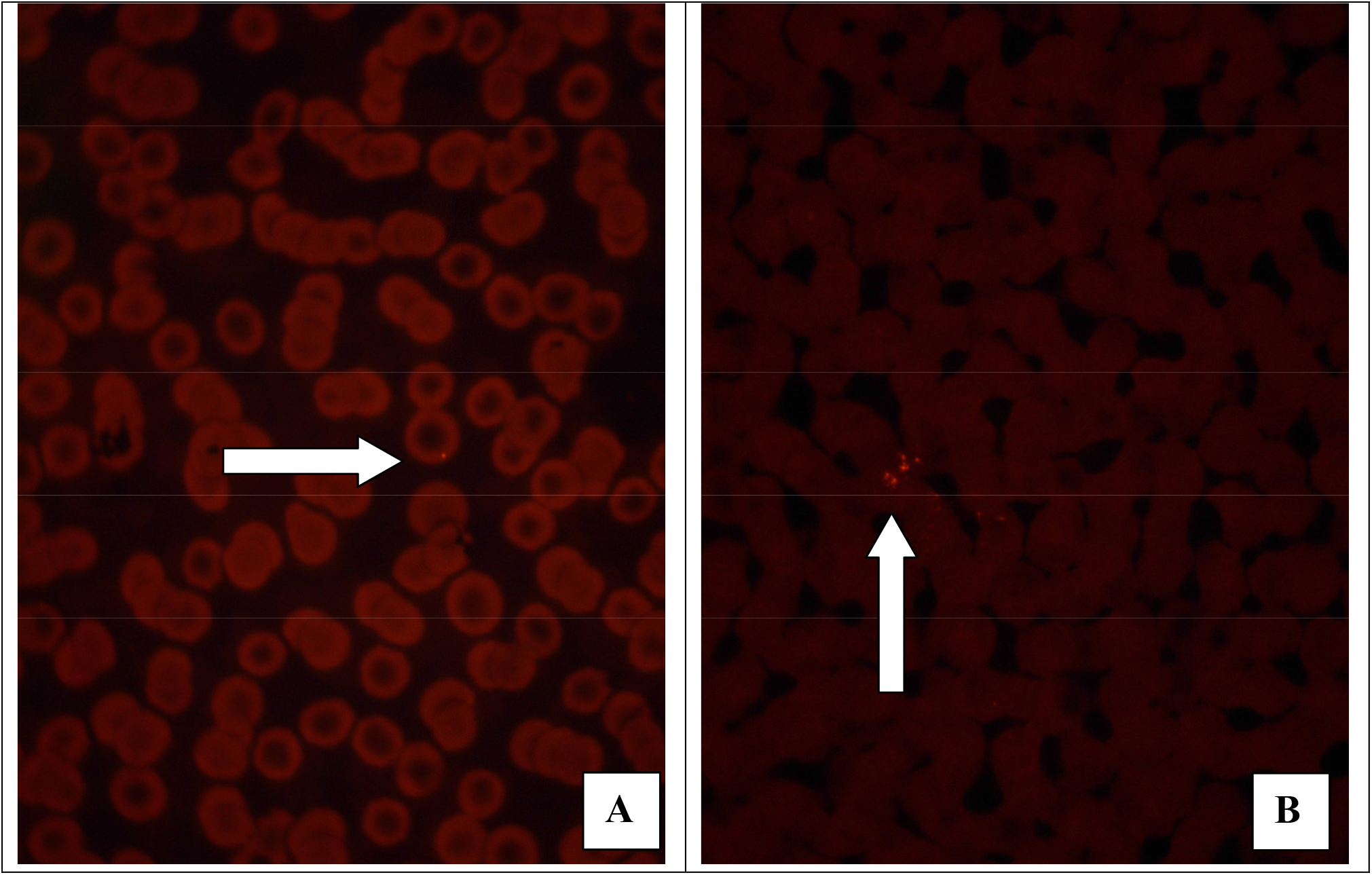
**FISH reaction to RNA SARS-Cov-2 in blood smears. Fluorescence microscopy. (A) - is a moderate course. Single foci of viral RNA hybridization in single red blood cells (shown by arrow). (B) - is a severe course of a new coronavirus infection. Moderate number of infected blood cells (shown by arrow). Multiplicity of magnification 10 * 100, immersion.**

**Fig. 3.**
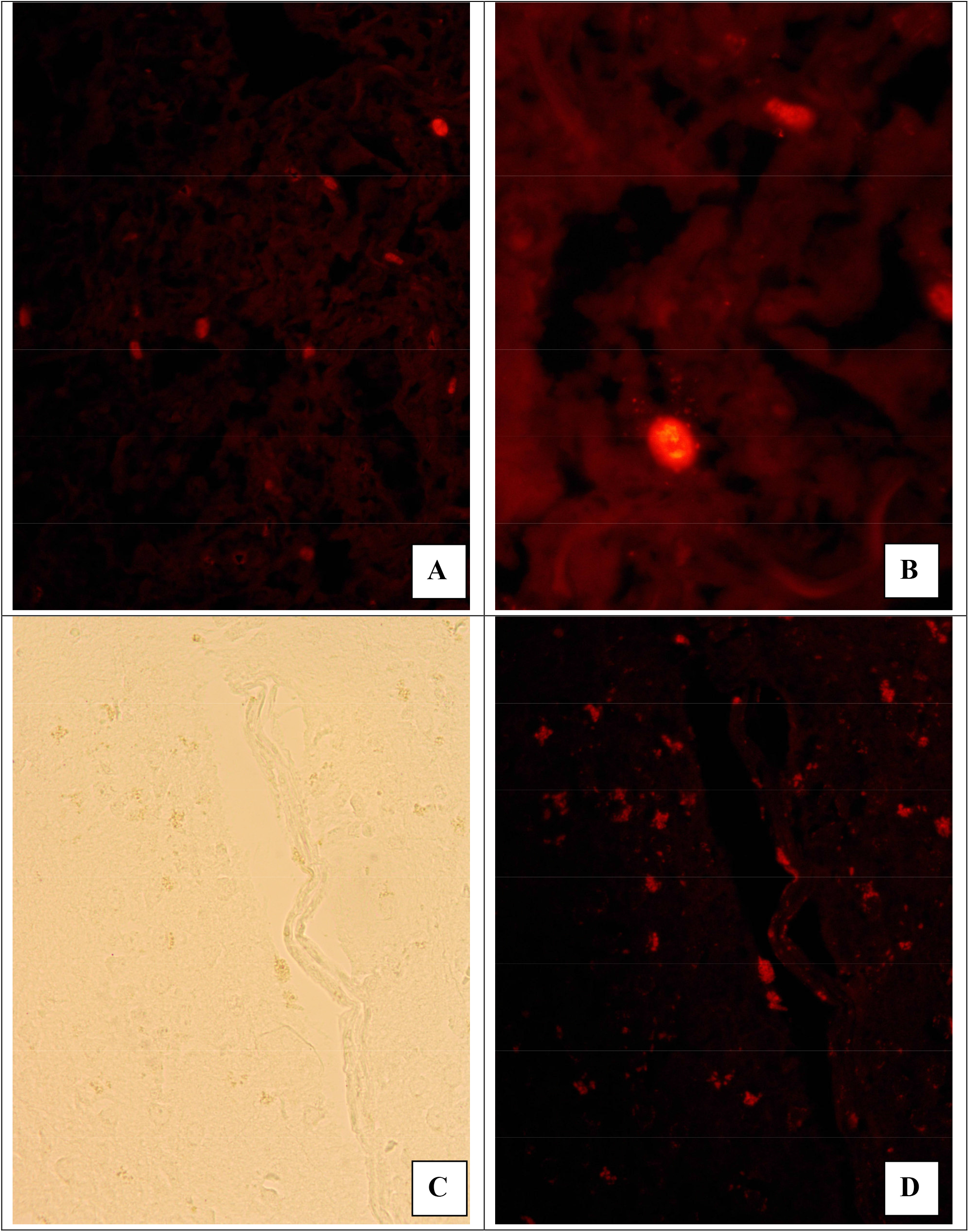
**FISH to RNA SARS-Cov-2 in lung and brain tissues (autopsy material). Fluorescence microscopy. (A) - Numerous foci of viral particle replication in the lungs. Magnification 10*40. (B) - Lung tissue. Viral RNA particles in the nucleus and cytoplasm of the cell (presumably alveolar macrophage). Multiplicity of magnification 10*100, immersion. (C) - brain tissue, multiple foci of lumpy brain decay. Light microscopy. (D) - is the same micropreparation. Fluorescence of hybridized viral RNA in these same destroyed cells. Multiplicity of magnification. 10*40.**

The highest expression of viral RNA was observed in the tissues of the internal organs of individuals who died from NCI. Studies of lung tissue and cerebral cortex are presented as a demonstration. In the control group, there was no fluorescence of ASO-TAMRA markers, while in the event of death from NCI, multiple fluorescence of markers in lung cells was detected (Fig. 3A). At a magnification of 10*100, it was seen that hybridization of ASO-TAMRA markers was both in the nucleus and in the cytoplasm (Fig. 3B). In the cortical tissue of the brain, light microscopy revealed multiple foci of block decay of neurons and glia, shown in Fig. 3C, visible even on an unstained hematoxylin and eosin section. When examining this site at fluorescence, it was found that the dead cells showed high saturation of viral RNA, which gave hybridization with the ASO-TAMRA fluorescent marker (Fig. 3D).

## DISCUSSION OF THE OBTAINED RESULTS

In the study, it was shown that ASO of the 5’-AGC CGA GTG ACA GCC ACA CAG type to the SARS-Cov-2 trs-gene region has a high binding specificity and a unique nucleotide sequence. The correct choice *in silico* of the nucleotide sequence of the blocking region of the gene and its complementary ASO is a guarantee of specificity and the absence of hybridization-dependent toxicity due to non-specific hybridization with other single-stranded nucleic acids in the body (12). The choice of the trs-gene is due to its high conservatism in coronaviruses. The low level of mutations of this gene determines its suitability for inhibiting the transcription of the entire coronavirus genome, which is proved by the effective suppression of intracellular reproduction of the virus in the plaque inhibition reaction in the present study. In therapeutic and prophylactic use in cell culture, the value of the effective dosage of ED_50_, which reduces the virus titer by 50%, was 3.84 mcg/ml (0.000384% solution). Considering the absence of toxicity at dosages up to 100 mcg/ml, the chemotherapeutic index is greater than 28.8. These results allow us to recommend this ASO for further research as a promising drug for NCI. Taking into account the conservatism of the SARS-Cov-2 trs-gene, the studied ASO can also be used for diagnostic studies in the fluorescence in situ hybridization reaction. The study of ASO conjugated with the TAMRA dye showed that this fluorochrome marker can be used in the diagnosis of SARS-Cov-2 in the FISH reaction on cytological preparations of nasopharyngeal and blood smears, as well as on histological preparations of deceased patients. Detection of viral RNA in the FISH reaction with ASO-TAMRA can visualize the presence of viral RNA in the nasopharyngeal discharge even in mild NCI. At the same time, when examining blood smears of patients with NCI, it was found that the detection of viral RNA begins only in moderate and severe patients, which is associated with the penetration of the virus into the systemic circulation. Therefore, the detection of a virus in the blood cells can serve as a diagnostic marker for the penetration of the virus outside the pulmonary system and the beginning of the generalization of the infection. In addition, intracellular detection of viral RNA using ASO-TAMRA in the absence of a clinical picture characteristic of HCI may be a sign of latent intracellular carriage of the virus.

The use of ASO-TAMRA is also effective for the detection of viral RNA in the tissues of internal organs. This method can be useful for determining the virus in clinical practice in intravital biopsy diagnostics and post-mortem diagnostics during autopsy, as well as for research purposes to determine the mechanisms of spread and localization of the SARS-Cov-2 virus.

## CONCLUSIONS

An antisense oligonucleotide of the 5’-AGC CGA GTG ACA GCC ACA CAG type, complementary to the trs-gene region of the SARS-Cov-2 coronavirus, is relevant for the available SARS-Cov-2 sequenced sequences and can be used for therapeutic and diagnostic purposes.

2. Antisense oligonucleotide to SARS-Cov-2 species 5’-AGC CGA GTG ACA GCC ACA CAG with phosphorothioate modification of internucleoside phosphate and 2’-oxymethyl modification of pentose showed high antiviral activity against SARS-Cov-2 by assessing plaque inhibition. In therapeutic and prophylactic use in cell culture, the value of the effective dosage of ED_50_, which reduces the virus titer by 50%, was 3.84 mcg/ml (0.000384% solution). Considering the absence of toxicity at dosages up to 100 mcg/ml, the chemotherapeutic index is greater than 28.8.

3. An antisense oligonucleotide of the type 5’-AGC CGA GTG ACA GCC ACA CAG conjugated with a fluorescent marker of tetramethylrhodamine (TAMRA) can be used as a diagnostic DNA probe for the detection of SARS-Cov-2 virus RNA by fluorescent in situ hybridization on cytological preparations smears of the nasopharynx, blood smears and tissue preparations of internal organs.

## Notes

### Competing Interest Statement

The authors have declared no competing interest.

## REFERENCES

1. Clinical laboratory diagnostics: textbook / Edited by V.V. Dolgova, FGBOU DPO “Russian Medical Academy of Continuing Professional Education”. -M.: FGBOU DPO RMANPO, 2016. ISBN 978-5-7249-2608-9

2. COVID-19 Treatment Guidelines Panel. Coronavirus Disease 2019 (COVID-19) Treatment Guidelines. National Institutes of Health. Available at https://www.covid19treatmentguidelines.nih.gov/. Accessed [07.24.2022].

3. Goryachev A.N., Kalantarov S.A., Tkachev V.V., Severova A.G., Goryacheva A.S. Potencial’naya vozmozhnost’ antismyslovoj terapii COVID-19 // Sovremennye problemy nauki i obrazovaniya [Potential Opportunity of Antisense Therapy of COVID-19// Modern Problems of Science and Education] doi: 10.17513/spno.30269– 2020. – ? 6. URL: https://science-education.ru/ru/article/view?id=30269 (Accessed: 24.07.2022)

4. Jansen GJ, Wiersma M, van Wamel WJB, Wijnberg ID. Direct detection of SARS-CoV-2 antisense and sense genomic RNA in human saliva by semi-autonomous fluorescence in situ hybridization: A proxy for contagiousness? PLoS One. 2021 Aug 17;16(8):e0256378. doi: 10.1371/journal.pone.0256378. PMID: 34403446; PMCID: PMC8370601.

5. Korzhevsky D.E., Gilyarov A.V. [Fundamentals Principles of Histological Technique]. St. Petersburg: “SpetsLit Publishing House” Ltd.; 2010. 96 p.

6. Kula-Pacurar A, Wadas J, Suder A, Szczepanski A, Milewska A, Ochman M, Stacel T, Pyrc K. Visualization of SARS-CoV-2 using Immuno RNA-Fluorescence In Situ Hybridization. J Vis Exp. 2020 Dec 23;(166). doi: 10.3791/62067. PMID: 33427241.

7. Mironov A.N. editor. Guidelines for Preclinical Trials of Medicinal Products. Part 2. Moscow: Grif i K; 2012

8. Neuman B.W., Stein D.A., Kroeker A.D., Churchill M.J., Kim A.M., Kuhn P., Dawson P., Moulton H.M., Bestwick R.K., Iversen P.L., Buchmeier M.J. Inhibition, escape, and attenuated growth of severe acute respiratory syndrome coronavirus treated with antisense morpholino oligomers. Journal of Virology. 2005. no. 79. P. 9665–9676.

9. Park M, Won J, Choi BY, Lee CJ. Optimization of primer sets and detection protocols for SARS-CoV-2 of coronavirus disease 2019 (COVID-19) using PCR and real-time PCR. Exp Mol Med. 2020 Jun;52(6):963–977. doi: 10.1038/s12276-020-0452-7. Epub 2020 Jun 16. PMID: 32546849; PMCID: PMC7295692.

10. Pharmacology of Antisense Drugs C. Frank Bennett, Brenda F. Baker, Nguyen Pham, Eric Swayze, and Richard S. Geary Annual Review of Pharmacology and Toxicology. 2017. vol. 57. no. 1. P. 81-105.].

11. Saifitdinova A.F. Two-dimensional fluorescence microscopy for the analysis of biological samples. SPb.: 2011; 110 (in Russian)

12. von Stillfried S, Boor P. Nachweismethoden von SARS-CoV-2 in Gewebe [Methods of SARS-CoV-2 detection in tissue]. Pathologe. 2021 Mar;42(2):208-215. German. doi: 10.1007/s00292-021-00919-8. Epub 2021 Mar 1. PMID: 33646360; PMCID: PMC7919251.

